# Characterizing Low-cost Registration for Photographic Images to Computed Tomography

**DOI:** 10.1101/2023.09.22.558989

**Authors:** Michael E. Kim, Ho Hin Lee, Karthik Ramadass, Chenyu Gao, Katherine Van Schaik, Eric Tkaczyk, Jeffrey Spraggins, Daniel C. Moyer, Bennett A. Landman

**Affiliations:** Vanderbilt University, Department of Computer Science, Nashville, TN USA; Vanderbilt University, Department of Electrical Engineering, Nashville, TN, USA; Vanderbilt University Medical Center, Department of Radiology and Radiological Sciences, Nashville, TN, USA; Tennessee Valley Healthcare System, Department of Veterans Affairs, Nashville, TN, USA; Vanderbilt University Medical Center, Department of Dermatology, Nashville, TN, USA; Vanderbilt University, Department of Biomedical Engineering, Nashville, TN, USA; Vanderbilt University School of Medicine, Department of Cell and Developmental Biology, Nashville, TN, USA; Vanderbilt University Institute of Imaging Science, Nashville, TN, USA

**Keywords:** Surface Registration, Computed Tomography, Photogrammetry, NeRF

## Abstract

Mapping information from photographic images to volumetric medical imaging scans is essential for linking spaces with physical environments, such as in image-guided surgery. Current methods of accurate photographic image to computed tomography (CT) image mapping can be computationally intensive and/or require specialized hardware. For general purpose 3-D mapping of bulk specimens in histological processing, a cost-effective solution is necessary. Here, we compare the integration of a commercial 3-D camera and cell phone imaging with a surface registration pipeline. Using surgical implants and chuck-eye steak as phantom tests, we obtain 3-D CT reconstruction and sets of photographic images from two sources: Canfield Imaging’s H1 camera and an iPhone 14 Pro. We perform surface reconstruction from the photographic images using commercial tools and open-source code for Neural Radiance Fields (NeRF) respectively. We complete surface registration of the reconstructed surfaces with the iterative closest point (ICP) method. Manually placed landmarks were identified at three locations on each of the surfaces. Registration of the Canfield surfaces for three objects yields landmark distance errors of 1.747, 3.932, and 1.692 mm, while registration of the respective iPhone camera surfaces yields errors of 1.222, 2.061, and 5.155 mm. Photographic imaging of an organ sample prior to tissue sectioning provides a low-cost alternative to establish correspondence between histological samples and 3-D anatomical samples.

## 1. INTRODUCTION

Integrating physical space information with volumetric information is essential for localizing histological data, which is acquired at the micron-scale, within the broader anatomical context of the human body, acquired at the millimetric-scale for organs and the centimetric-scale for systems. The Human BioMolecular Atlas Program (HuBMAP) is working to map the complete human body at the cellular level in a way that is generalizable across populations (https://hubmapconsortium.org/)^1^. HuBMAP teams are generating extensive 3D templates of the human body along with rich molecular and cellular data, so techniques to localize small tissue biopsies and blocks to larger anatomical features and organ systems are of increasing immediate importance.

Computed Tomography (CT) and Magnetic Resonance Imaging (MRI) are imaging modalities widely used to obtain structural information of small anatomical structures in vivo for diagnosis and preoperative planning due to the high-resolution volumetric 3-D images they produce^2–8^. Several 3-D CT HuBMAP atlases, such as the kidney with substructure and eye atlases from Lee et al.^9–11^, the pancreas atlas from Zhou et al. ^12^, and the gut cell atlas ^13,14^ have been released in an organ specific manner. These data frames are being brought together within an overall systems perspective to help generalize structural, cellular, and molecular biomarkers across populations and different morphologies. Yet, routine, automated localization is not performed to identify the location of samples within the overall coordinate systems.

Meanwhile, 3-D computer vision techniques are widely used in the reconstruction of surface information from photogrammetry (e.g. stereoscopic techniques). Krishna et al. use stereoscopic scanning electron microscope images to construct surface topography for characterization of engineering surface quality^15^. HajiRassouliha, et al. use a four-camera stereoscope to measure 3-D deformations of skin^16^. Another common technique for 3-D surface construction from photogrammetry is structure from motion (SfM), in which 3-D structure is inferred from 2-D imaging sequences. Su et al. use digital surface models using photogrammetry from unmanned aerial vehicles to phenotype corn crops based on height^17^. Um et al. examine 3-D surface reconstruction using microscopic SfM^18^. Shilov et al. use SfM to construct surfaces of feet^19^. Zhan et al. propose a method for registration of pointclouds obtained from photogrammetric data and CT of a directional gyro^20^. However, methods for obtaining anatomically accurate surfaces of soft tissue with sub-millimetric precision require expensive or specialized hardware, such as Holocreators (https://holocreators.com/) and 3Dmd (https://3dmd.com/) hardware.

Despite these reported methods and the existence of these initial HuBMAP atlases, there needs to be a common procedure for this mapping of multimodal data into volumetric atlases. Given that the data acquisition sites of the HuBMAP project are diverse and widespread, simpler methods of multimodal data integration are more realistic options compared to more expensive or complex ones that require specialized expertise or equipment. Thus, there is a need to identify and establish both a technology and procedure that can fit within such a dynamic, multi-site project.

In this paper, we characterize the registration of surfaces generated using two methods low-cost photogrammetry to surfaces from CT scans. One method uses a commercial camera that creates surfaces from photogrammetry with a binocular lens, while the other uses a common smartphone video that is subsampled for surface reconstruction of a scene. We characterize the quality of these surfaces through registration of the photogrammetric surfaces to surfaces obtained from CT scans (Figure 1).

**Figure 1.**
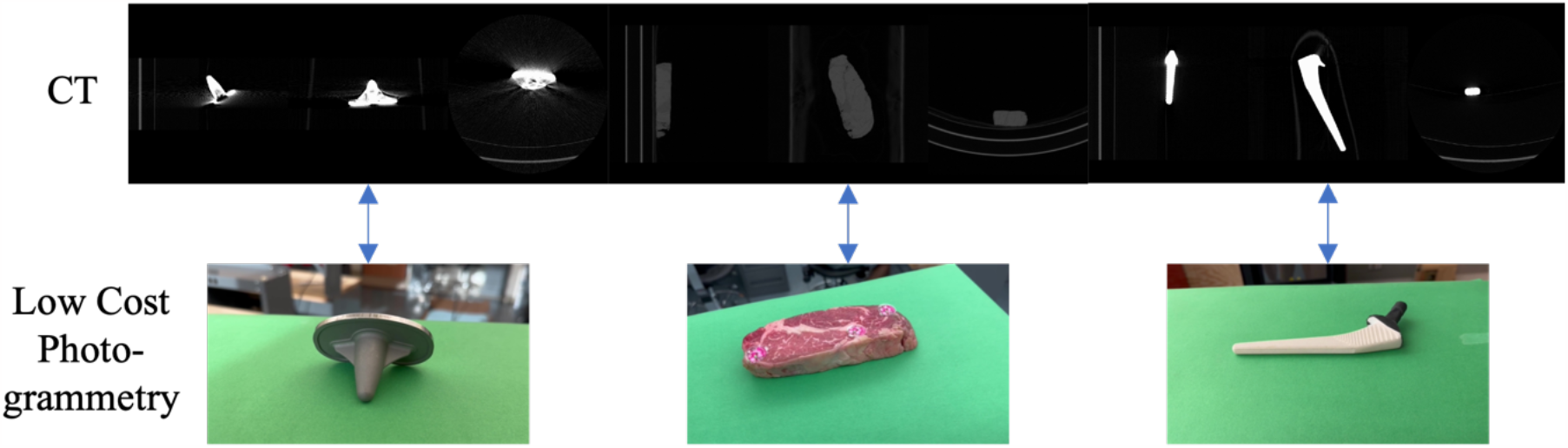
We map surfaces of phantom objects obtained from low-cost photogrammetry to surfaces obtained from volumetric CT scans of the phantoms in order to examine if similar techniques could be applied to mapping of histology data to volumetric CT data. Knee (left) and hip (right) implants are static objects with reflective and non-reflective surfaces respectively whose surfaces should be easily mappable. Chuck-eye steak (center) is a moderately deformable phantom that also provides a slightly reflective surface, which should be more similar to biological tissue.

## 2. METHODS

CT images of a hydroxyapatite stem of a CORAIL hip implant from DePuy Synthes with black non-reflective tape covering the metal head, a tibial tray of a P.C.F. SIGMA RP knee replacement from DePuy Synthes, and a chuck-eye steak from Bare Bones Butcher in Nashville, TN were acquired with a 2017 Philips Vereos PET/CT system with integrated 64-slice CT scanner. Scanning of the hip implant was at a resolution of 0.325521 x 0.325521 x 0.669983 mm^3^, the steak at 0.626953 x 0.626953 x 1 mm^3^, and the knee implant at 0.325521 x 0.325521 x 0.670013 mm^3^. 1.5mm Beekley Medical X-SPOT mammography skin markers were placed on the steak to identify landmarks for registration accuracy. Knee and hip implant landmarks were selected based on structural points of interest. Triangulated meshes were generated from segmentations of the CT images using nii2mesh (https://github.com/neurolabusc/nii2mesh.git). Surfaces for the steak were obtained with and without CT markers. The steak surface with CT markers was only used to identify fiducial location on the surface.

Triangulated meshes of the three objects were obtained from photogrammetry using two different methods. One method is via the general capture feature of the VECTRA® H1 handheld imaging system developed by Canfield Technologies (https://www.canfieldsci.com/imaging-systems/vectra-h1-3d-imaging-system/), which uses a specialized mirror lens to take a single binocular picture of an object to create a mesh with millimetric calibration of object dimensions. De Stefani et al. examine the validation of VECTRA systems for 3-D facial reconstruction in a review paper but did not examine accuracy of the general capture feature^21^. Another method involves the synthesis of a scene from photogrammetry using Neural Radiance Fields (NeRF)^22^. 1080p resolution videos of varying length (Table 1) were taken on an iPhone 14 Pro and subsampled at four frames per second to create a sequential batch of images. Position from motion of the pictures was calculated using COLMAP^23,24^ with sequential image matching, meaning that images were paired based on sequential proximity to each other. Positional information was input along with the images into instant-npg software to generate the scene^25^ using an axis-aligned bounding box (aabb) scale of 4.

**Table 1.** Lengths of videos for NeRF surfaces. An asterisk (*) indicates videos that produced surfaces not used in the analysis.

| Object | Video 1 (s) | Video 2 (s) | Video 3 (s) | Video 4 (s) | Video 5 (s) |
| --- | --- | --- | --- | --- | --- |
| Hip Implant | 45 | 37* | 35 | 35 | 40 |
| Knee Implant | 45 | 42 | 44 | 36 | 32* |
| Chuck-eye Steak | 36* | 78 | 60 | 47 | 49 |

Photogrammetry was obtained in a roughly 15 x 28 ft room with ambient and fluorescent lighting present in an area that was roughly 12 x 8 ft and free of objects. Photogrammetry for the H1 camera was obtained using sky blue Yizhily seamless photography photo backdrop paper as a background. Objects were photographed on the floor of the room on top of the backdrop paper (Figure 2). Hip and knee implants were placed on a metal rod 19.5 in tall to introduce parallax between the object and the background. Steak was placed on the background paper with a layer of plastic wrap underneath the steak. Surfaces were output in OBJ format with a texture file that referenced a PNG file for color information. Python (v3.10.11) code was implemented to convert the OBJ and PNG files to a visualization toolkit (VTK) surface file.

**Figure 2.**
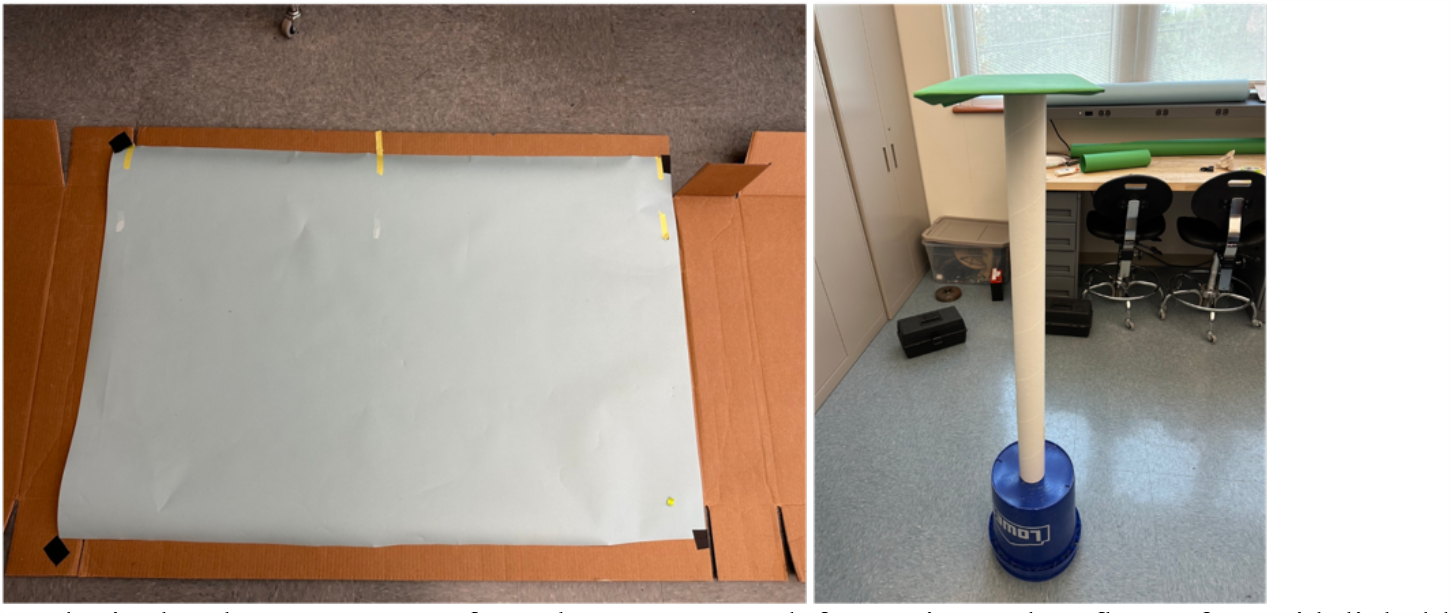
Setup to obtain the photogrammetry from the H1 camera (left) requires only a flat surface with light blue background paper and enough space to take a photo with the camera. Setup for the iPhone videos (right) uses a green background platform underneath the object with enough space to circle around the object completely, although any color background paper can be used as long as it is easily differentiable from the surface of interest and not very reflective. The platform was raised to allow for easier movement around the setup while holding the iPhone. Both setups use only ambient and fluorescent lighting, were free of nearby clutter and objects, and are low-cost with minimal preparation.

Photogrammetry for NeRF surfaces was taken with objects placed directly on a 10 x 17.5 in piece of cardboard that was covered with the stinger (green) Yizhily seamless photography photo backdrop paper. The cardboard surface was placed on top of a 43 in cardboard tube and a 14 in bucket to place the object at roughly eye level for easier acquisition (Figure 2). Surfaces were generated using the instant-npg interactive GUI to crop out as much of the scene as possible while keeping the entirety of the object in the scene and were exported in PLY format that was converted to VTK format.

Both batches of surfaces were preprocessed using the VTK^26^ and trimesh (v3.22.0) (https://github.com/mikedh/trimesh) libraries to remove points on the output meshes that were not part of the object surface. H1 surfaces were thresholded to remove points that were blue in color and retained the connected component that contained the object. NeRF surfaces were thresholded to remove points that were green in color and keep the connected component containing the object.

### 2.1 Surface Registration

Prior to registration, both the static and moving surfaces were first centroid centered using python code. Initial alignment of centroid-centered surfaces was performed via rigid registration of the principal component axes of the surfaces with python. These alignments were visually inspected for quality of alignment. If the orientation was not proper, manual rigid rotation along the axes with python was applied to achieve a better alignment. NeRF surfaces required an additional degree of freedom for volumetric scaling as the surface dimensions are not calibrated to match the object that was scanned. H1 surfaces were calibrated millimetric surfaces and thus did not require an additional scaling. After preliminary alignment and scaling, the iterative closest point (ICP) algorithm ^27^ was used to rigidly register the surfaces using the torch3D library (https://github.com/facebookresearch/pytorch3d.git). NeRF surface registration using the torch3D ICP also used the scaling option to perform a second scaling of the mesh. To mitigate the effect that noise points from NeRF surfaces have on the registration process, the ICP algorithm was computed for registering the CT surface to the NeRF surface, and the inverse transformations were applied to the NeRF surfaces to move them into the same space as the CT surfaces. Figure 3 illustrates this process for both batches of surfaces.

**Figure 3.**
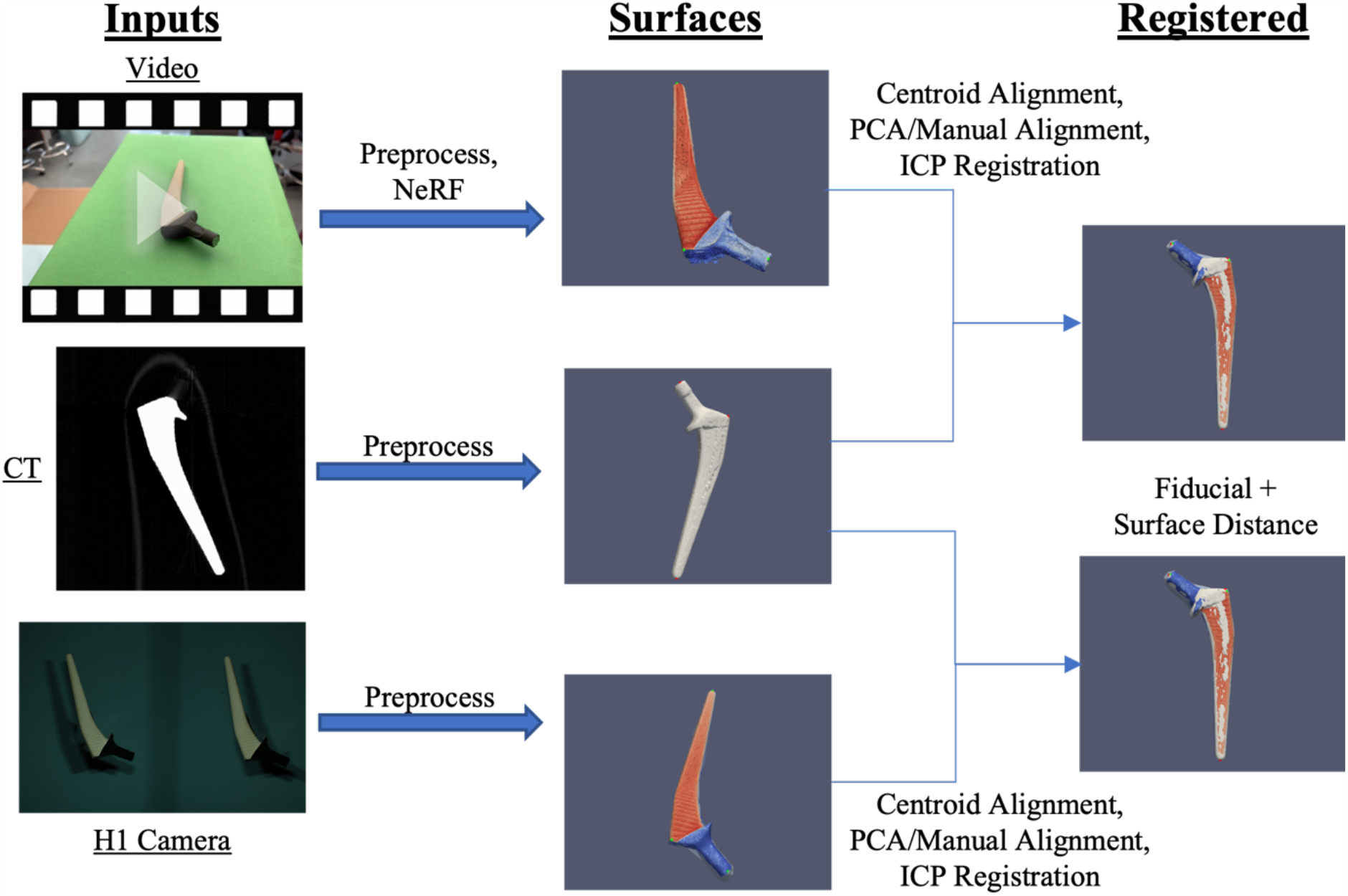
CT scans of objects were segmented and processed to obtain a 3-D surface. Photogrammetry surfaces of the same objects were obtained through subsampled 1080p video input from an iPhone 14 Pro and binocular images from a VECTRA H1 camera. After an initial centroid alignment and PCA alignment with manual inspection, the photogrammetry surfaces were registered via the ICP algorithm. Alignment of the surfaces was assessed from fiducial registration error and surface distances.

Registrations of both batches of surfaces are evaluated for goodness of fit with two metrics. First, by average fiducial distance error of each of the three fiducials on each object. Fiducials on the surfaces of the steak were identified by the CT markers, whereas fiducials on the hip and knee implants were selected based on landmark position. Second, by the minimum root mean squared error (RMSE) surface distance, calculated by squaring the distances from each vertex on the moving surface to the closest point on the target surface. Code for the preprocessing and registration can be found here: https://github.com/MASILab/SurfaceRegistration.git.

## 3. RESULTS

We characterize the registration error as the distance (mm) between respective fiducials after registration of the photogrammetry surfaces to the CT surface (Figure 4). We report the error of successful surface reconstructions and registrations. For each of the five videos we took of each object for NeRF surfaces, four successfully produced surfaces that could be registered to the respective CT surface. Videos that failed produced surfaces with too much noise to accurately capture the surface information of the object. One surface of the hip implant from the H1 camera did not fully capture the entire visible surface, and thus was not used in the analysis. We observe H1 camera surfaces for both the hip implant and the steak yielded a better on average registration accuracy for fiducial distance than the NeRF surfaces with a two to three times larger error for fiducial registration for the NeRF surface (Table 2). Conversely, we observe the NeRF surface of the knee implant yielded a closer registration than the H1 surface with a two to three times larger error.

**Table 2.** Average fiducial registration error and RMSE surface distance for photogrammetry surfaces and CT surface registration with one standard deviation of error.

| Surface<br>(n=26) | Mean<br>Fiducial 1<br>Error (mm) | Mean<br>Fiducial 2<br>Error (mm) | Mean<br>Fiducial 3<br>error (mm) | Average<br>Fiducial<br>Error (mm) | Average<br>RMSE<br>Surface<br>Distance<br>(mm) |
| --- | --- | --- | --- | --- | --- |
| Hip Implant<br>(H1) (n=4) | 2.063 +/- 0.309 | 1.691 +/- 0.419 | 1.486 +/- 0.389 | 1.747 +/- 0.445 | 0.616 +/- 0.113 |
| Knee Implant<br>(H1) (n=5) | 3.077 +/- 1.673 | 4.211 +/- 1.588 | 4.507 +/- 2.667 | 3.932 +/- 2.127 | 1.370 +/- 0.433 |
| Chuck-eye<br>Steak (H1)<br>(n=5) | 1.102 +/- 0.242 | 1.629 +/- 0.426 | 2.346 +/- 0.453 | 1.692 +/- 0.639 | 0.671 +/- 0.038 |
| Hip Implant<br>(NeRF) (n=4) | 1.619 +/- 0.497 | 1.138 +/- 0.776 | 0.909 +/- 0.316 | 1.222 +/- 0.635 | 2.067 +/- 0.711 |
| Knee Implant<br>(NeRF) (n=4) | 2.313 +/- 0.681 | 2.420 +/- 1.174 | 1.449 +/- 0.337 | 2.061 +/- 0.917 | 3.073 +/- 0.693 |
| Chuck-eye<br>Steak (NeRF)<br>(n=4) | 5.919 +/- 2.124 | 4.063 +/- 1.665 | 5.483 +/- 1.961 | 5.155 +/- 2.083 | 6.151 +/- 0.501 |

**Figure 4.**
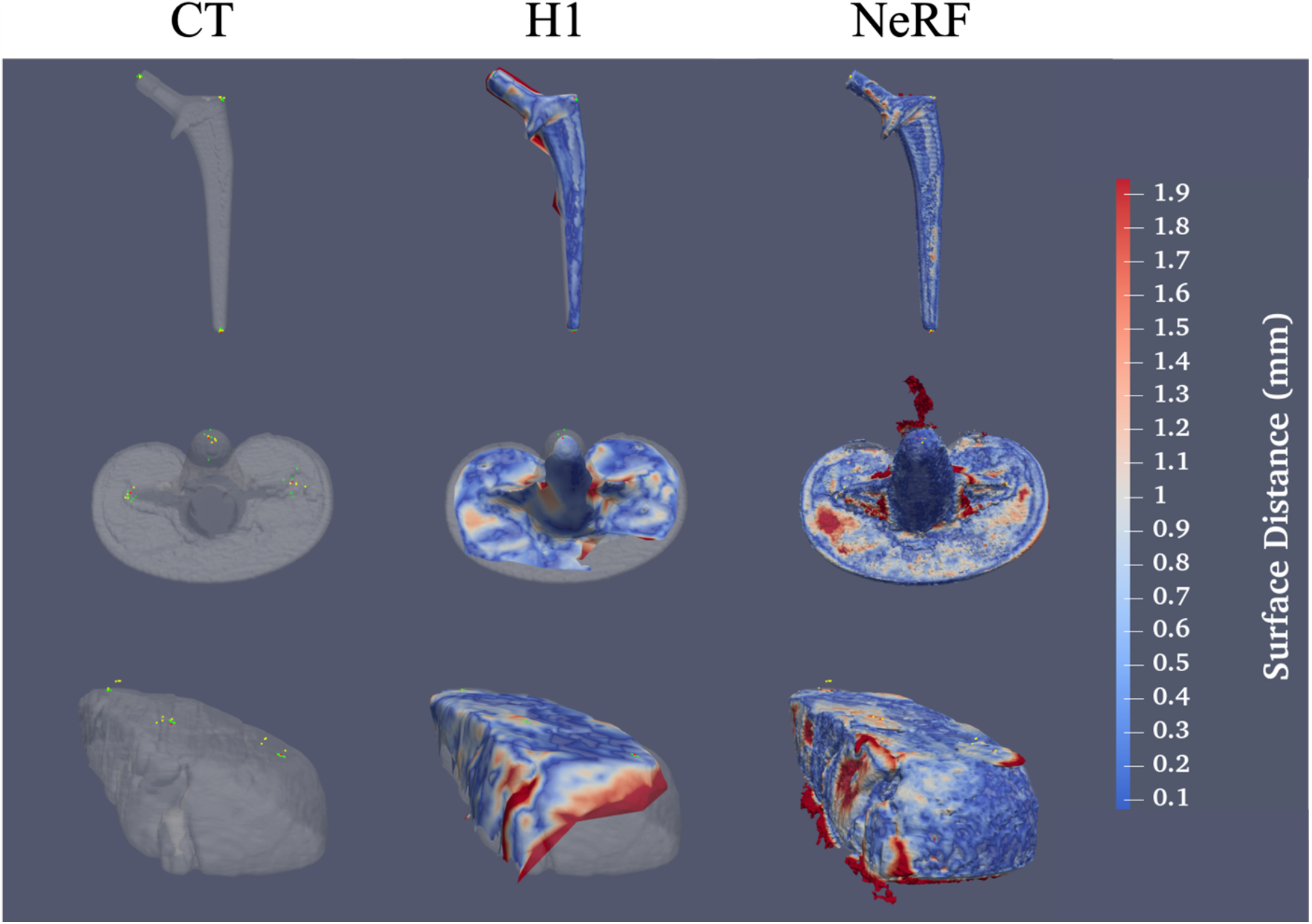
NeRF surfaces capture much finer detail and sharper edges than surfaces from the VECTRA H1 camera but are much noisier and more susceptible to artifacts from light reflections. (Top) From left to right, a surface obtained from a CT of a hip implant with fiducials from the CT surface and both H1 and NeRF surface fiducials overlaid, an H1 surface of a hip implant registered to the CT surface, and a NeRF surface registered to the CT surface. Red markers indicate CT fiducials, green indicate H1 surface fiducials, and yellow indicate NeRF surface fiducials. Similar results are shown for the knee implant (center) and the chuck-eye steak (bottom). Surface colors indicate the surface distance of each point on the photogrammetry surface to the CT surface.

We observe that, for all 3 objects, the average surface distance for the NERF surfaces is greater than that of the H1 camera surfaces. We attribute a large amount of this error to artifactual points on the interior of the surface synthesized by the NeRF, as surface information should not contain any information about the interior of the object. Thus, average MSE surface distance may not be representative of the registration accuracy.

## 4. DISCUSSION

Results from Table 2 suggest that the optimal method for obtaining low-cost photogrammetry surfaces is context dependent: objects that have sharp changes in depth or non-smooth surfaces may yield NeRF meshes that are more morphologically accurate than H1 camera meshes. However, for objects that are smoother and more uniform, results suggest the H1 camera yields more accurate surfaces. Additional error in fiducial distance for registration of NeRF surfaces may be attributable to the CT markers used to mark fiducial location. The markers were not included in the registration process for the CT surface; however, their structure was visible on the NeRF surfaces as opposed to the H1 surfaces on which their structure was not visible (Figure 4). This additional distance likely resulted in the fiducial distances for the NeRF surfaces to be ∼1mm off the true fiducial distance. Surface meshes obtained from NeRF appear to be more prone to have artifactual information from light reflections, which is evidenced by the non-uniform color of the surfaces as compared to the objects themselves as well as noise points that can be found coming off of the surfaces (Figure 4). Such errors could have attributed to the increased fiducial error in NeRF surfaces as opposed to the knee and hip implants (Table 2).

H1 Camera surfaces have relatively high accuracy, making them a candidate method for the HuBMAP processing protocol. Fiducial error on the NeRF surfaces suggest that the methodology employed in this work is not yet suitable for mapping histological information to CT data. However, further refinement of the surfaces could yield more promising results. Nonetheless, acquisitions for both NeRF and H1 surfaces are simple, low-cost, and can be easily integrated into the data acquisition pipeline for HuBMAP. System design requirements include an iPhone 14 Pro (currently the latest model) for NeRF or the VECTRA H1 camera for the other, both of which are handheld. We note that at the time this work was completed, Canfield has come out with a newer model, the H2 camera, for the VECTRA handheld 3-D imaging line. Setup for acquisition would only require low-cost, non-reflective, colored background paper without any specialized lighting, as well as sufficient distance to image the object from or enough space to move around the object for a video. Acquisition of the videos and images themselves takes only a few minutes at most. Thus, we believe a methodology similar to ours would be easy to integrate into data acquisition for projects such as HuBMAP, following a characterization of these techniques with biological specimens.

While we only examined the NeRF surfaces generated from a fixed frame rate and aabb value, these parameters could be optimized to generate more accurate scenes and thus more morphologically accurate meshes. Videos captured of objects were acquired at 29.99 frames per second but were only sampled at four frames per second to create sets of sequential images. Future work could examine whether an increased sampling rate would increase or decrease the amount of noise and accuracy in the surface of objects. We also did not apply any smoothing filters to NeRF meshes. Applying smoothing algorithms similar to the one described in Shilov et al. for smoothing 3-D surfaces of feet could increase the quality of registration^19^.

Another potential future direction for this work includes exploring deformable registration of photogrammetric surface information from human organ tissue to CT surface information. Organ tissue is much more deformable than static objects or muscle tissue that was explored in this paper, and such advancements could lay the groundwork for mapping photogrammetry to organs *in vivo*.

## ACKNOWLEDGEMENTS

This research is supported by the NIH Common Fund U54DK134302 and U54EY032442 (Spraggins), NSF CAREER 1452485, NIH 2R01EB006136, NIH 1R01EB017230 (Landman), IK2 CX001785 (Tkaczyk), and NIH R01NS09529. The Vanderbilt Institute for Clinical and Translational Research (VICTR) is funded by the National Center for Advancing Translational Sciences (NCATS) Clinical Translational Science Award (CTSA) Program, Award Number 5UL1TR002243-03. The content is solely the responsibility of the authors and does not necessarily represent the official views of the NIH.

